# Thermostable mutants of glycoside hydrolase family 6 cellobiohydrolase from the basidiomycete *Phanerochaete chrysosporium*

**DOI:** 10.1101/2020.05.11.089235

**Authors:** Sora Yamaguchi, Naoki Sunagawa, Mikako Tachioka, Kiyohiko Igarashi, Masahiro Samejima

## Abstract

Thermal inactivation of saccharifying enzymes is a crucial issue for the efficient utilization of cellulosic biomass as a renewable resource. Cellobiohydrolases (CBHs) is a kind of cellulase. In general, CBHs belonging to glycoside hydrolase (GH) family 6 (Cel6) act synergistically with CBHs of GH family 7 (Cel7) and other carbohydrate-active enzymes during the degradation of cellulosic biomass. However, while the catalytic rate of enzymes generally becomes faster at higher temperatures, Cel6 CBHs are inactivated at lower temperatures than Cel7 CBHs, and this represents a limiting factor for industrial utilization. In this study, we produced a series of mutants of the glycoside hydrolase family 6 cellobiohydrolase *Pc*Cel6A from the fungus *Phanerochaete chrysosporium*, and compared their thermal stability. Eight mutants from a random mutagenesis library and one rationally designed mutant were selected as candidate thermostable mutants and produced by heterologous expression in the yeast *Pichia pastoris*. Comparison of the hydrolytic activities at 50 and 60 ^°^C indicated that the thermal stability of *Pc*Cel6A is influenced by the number and position of cysteine residues that are not involved in disulfide bonds.

## INTRODUCTION

Cellulosic biomass is the most abundant carbon stock in nature, and its degradation to soluble sugars has the potential to replace fossil resources by providing an alternative raw material for fuels and chemicals. Further, as enzymatic saccharification of cellulosic biomass would not require strong acid or alkali, or intense heat, it should involve lower energy consumption than chemical or physical treatments. Cellobiohydrolases (CBHs) are a kind of cellulase and indispensable for the complete enzymatic hydrolysis of cellulose because they can degrade crystalline regions that are resistant to enzymatic hydrolysis by removing cellobiosyl units from the cellulose chain ends.^1)^ Nevertheless, the degradation of highly crystalline cellulose remains a key bottleneck for achieving efficient enzymatic saccharification.

CBHs that degrade crystalline cellulose are classified into either glycoside hydrolase family 6 (Cel6, EC.3.2.1.91) or 7 (Cel7, EC.3.2.1.176) in the Carbohydrate-Active enZyme (CAZy) database (http://www.cazy.org/).^2)^ Most Cel6 and Cel7 CBHs have a carbohydrate-binding module (CBM) and a catalytic domain (CD), which are connected by a flexible linker. Fungal Cel6 CBHs are important, because their hydrolytic activity is comparable to that of Cel7 CBHs, and synergistic hydrolysis occurs when crystalline cellulose is incubated with both Cel6 and Cel7 together.^3)4)^ However, the thermal stability of Cel6 CBHs is generally lower than that of Cel7 CBHs. For example, the optimum temperatures of Cel6 and Cel7 CBHs from the thermophilic filamentous fungus *Chaetomium thermophilum* are 50 ^°^C ^5)^ and 65 ^°^C,^1)^ respectively. This is a problem, because industrial-scale enzymatic saccharification is conducted at an elevated temperature to increase the hydrolysis rate. Therefore, increasing the thermal stability of Cel6 CBH should immediately lead to an increase in the efficiency of the commercial process.

Numerous studies have attempted to improve the thermal stability of Cel6 by applying two major strategies, i.e., random mutagenesis and rational design. Random mutagenesis is generally employed when information about the target enzyme is limited. On the other hand, in the process of rational design, key amino acid(s) to be changed are firstly identified based on the enzyme structure and the interaction between enzyme and substrate, and then the designed mutants are prepared and characterized.^6)^

The methylotrophic yeast *Pichia pastoris* is a suitable host for the expression of fungal Cel6 CBHs because it performs post-translational modifications found in the eukaryote and it can secrete the fungal proteins in up to gram quantities per liter of culture.^7)^ For example, production of Cel6A CBH from the wood-decaying fungus *Phanerochaete chrysosporium* (*Pc*Cel6A) in *P. pastoris* is as much as 4.6 g/L at 160 hours of cultivation.^8)^ Therefore, random mutagenesis combined with expression in *P. pastoris* has been employed to improve the catalytic efficiency and thermal stability of fungal Cel6 CBHs.^5)9)^ However, the contribution of each individual mutation to the activity of mutants with multiple mutations has not been investigated, e.g., by preparing mutants with each single mutation, although this information would be useful for rationally enhancing the activity even further.

Regarding the rational design of thermostable Cel6 CBHs, free cysteine (cysteine residues that do not form a disulfide bond) has been a target of substitution.^10)11)^ These studies analyzed the thermal stability of the mutants by measuring the incubation temperature at which the enzyme loses 50% of its activity, the residual activity, and the half-life. However, these methods do not provide information about the hydrolytic activity during incubation with substrates at elevated temperatures, though this ability is critical for achieving more efficient saccharification of cellulosic biomass.

In the present study, therefore, we heterologously expressed in *P. pastoris* a series of mutants of fungal *Pc*Cel6A with substitutions based on either random mutagenesis ^9)^ or rational design.^11)^ We compared the activities of these mutants by incubating them with amorphous phosphoric acid-swollen cellulose or crystalline cellulose III_I_ at different temperatures. Based on the results, we discuss the critical features for increased thermal stability of *Pc*Cel6A.

## MATERIALS AND METHODS

### Materials

DNA polymerases PrimeSTAR Max (TaKaRa Bio Inc., Shiga, Japan) and KOD-Plus (Ver.2; Toyobo Co., Ltd, Osaka, Japan) were used to amplify mutated DNA. One Shot^®^ TOP10 Chemically Competent *E. coli* (Thermo Fisher Scientific Inc., MA, USA) was used to amplify the plasmid. *Pme*I (New England Biolabs, MA, USA) was used to linearize the amplified plasmid for the transformation of *P. pastoris* strain KM71H, which was used for heterologous production of the mutant enzymes. Yeast (BD Biosciences, Miami, USA), peptone (Nihon Pharmaceutical Co., Ltd, Tokyo, Japan), and glycerol (FUJIFILM Wako Pure Chemical Corporation, Osaka, Japan) were used for the medium. Phosphoric acid-swollen cellulose (PASC) was prepared as reported.^12)^ Cellulose I_α_ and III_I_ were prepared from green algae *Cladophora spp*. according to the reported method.^13)^ *Aspergillus niger* β-glucosidase was acquired from Megazyme Ltd. (Wicklow, Ireland)

### Construction of PcCel6A expression plasmids

Primers for the site-directed mutations are listed in Table S1. Mutant libraries were constructed by inverse PCR using these primers, and pPICZα vector (Thermo Fisher Scientific) containing *Pc*Cel6A C240S/C393S gene with a *P. pastoris* codon bias was synthesized by Genscript Biotech Corporation (NJ, USA). PCR reaction mixture was purified with the Wizard^®^ SV Gel PCR Clean-Up System (Promega Corporation, WI, USA). One Shot^®^ TOP10 Chemically Competent *Escherichia coli* (Thermo Fisher Scientific) cells were transformed to amplify the mutated genes, and the plasmids were extracted from *E. coli*. Approximately 5 μg of each *Pc*Cel6A mutant gene in pPICZα was linearized by restriction enzyme *Pme*I (New England Biolabs) for the transformation of *P. pastoris* by electroporation.

### Enzyme expression in *Pichia pastoris*

The mutant *Pc*Cel6A library was produced by heterologous expression in *P. pastoris* strain KM71H (Thermo Fisher Scientific). *P. pastoris* containing wild-type *Pc*Cel6A gene was produced according to the reported method.^14)^ Colonies containing WT or mutant *Pc*Cel6A genes were grown in 1% yeast, 2% peptone (YP) medium with 2% glycerol at 30 ^°^C. Then, expression was induced in YP medium at 26.5 ^°^C by the addition of methanol (1% (*v/v*), final concentration) every other day. Aliquots of 80 μl of yeast culture were sampled for 3 days after the induction of the enzyme expression and centrifuged at 4 ^°^C with 10,000 x *g* for 10 min. The third-day culture was centrifuged at 4 ^°^C with 3,000 x *g* for 5 min and centrifuged again at 15,000 x *g* for 10 min at 4 ^°^C. The protein concentration of the supernatants was determined using Bio-Rad Protein Assay Dye Reagent Concentrate (Bio-rad Laboratories, Inc., CA, USA); the absorbance was measured at 595 nm with a Thermo Scientific Multiscan^®^ GO (Thermo Fisher Scientific). The expression level of enzymes was evaluated by SDS-PAGE using 12% polyacrylamide gel. A picture of the gel was taken with a CanoScan (Canon Inc., Tokyo, Japan) at 50% exposure time. The image was modified to change the shape of the lanes from trapezoidal to rectangular by using GIMP (Ver. 2.10.14.0, https://www.gimp.org) and converted to a 32-bit gray scale with ImageJ (Ver.1.52a, https://imagej.nih.gov/ij/). The amount of *Pc*Cel6A was estimated as the peak area of bands between 50 and 75 kDa.

### Activity measurements of crude enzymes

Culture supernatants of WT or mutant *Pc*Cel6A (10 μL) were incubated with 100 μg of PASC or cellulose III_I_ prepared as described previously^12)13)^ in 100 mM sodium acetate buffer (pH 5.0) using 96-well plates at 50 or 60 ^°^C with shaking at 1,000 rpm. After two hours of incubation, the solutions were filtered using 96-well plates with a 0.22 μm filter (MultiScreen^®^ Filter Plates, Merck Millipore, MA, USA). Then, 5 μL of 40 U/mL *Aspergillus niger* β-glucosidase (Megazyme Ltd.) was added to 100 μL of filtrate, and the plates were incubated at 60 °C for 48 h with shaking at 1,000 rpm. After hydrolysis, the solutions were heated at 98 ^°^C for 3 min and filtered again. The concentration of glucose in filtrate was quantified using Glucose CII-Test Wako (FUJIFILM Wako Pure Chemical Corporation) by measuring the absorbance at 492 nm with a Thermo Scientific Multiscan^®^ FC (Thermo Fisher Scientific).

## RESULTS AND DISCUSSION

Improving the thermal stability of fungal Cel6 CBHs is critical to increase the saccharification rate of cellulosic biomass, because these enzymes are the least stable among the cocktail of cellulolytic enzymes. The current study provides insights into the factors that determine the thermal stability of *Pc*Cel6A.

### Preparation of *Pc*Cel6A mutants

Amino acid residues that were expected to influence the thermostability of *Pc*Cel6A were chosen based on reported experimental results of random mutagenesis ^9)^ or rational design,^11)^ as shown in Fig. 1. Eight single mutations with different characteristics were chosen from the mutant library generated by introducing random mutations into wild-type *Pc*Cel6A in a *Pichia* expression system.^9)^ Cys25 is located on the CBM and forms a disulfide bond with Cys8 in the CBM. Ala103 and Ala105 are located at the surface of the CD. Met257 was proposed to stabilize the side chain of the adjacent residue.^15)^ Trp267 is located in the entrance of the active site tunnel. Gly346 and Gly421 lie within the loop near the active site. G421A was selected because G421D was expected to have a higher specific activity based on a previous study,^9)^ but the aspartic acid residue is large and acidic and might drastically change the enzyme structure. Cys240 and Cys393 should not form a disulfide bond because they are distant from other cysteine residues. The double mutant C240S/C393S was rationally designed as a candidate thermostable mutant by substituting the two cysteine residues with serine.

**Fig. 1.**
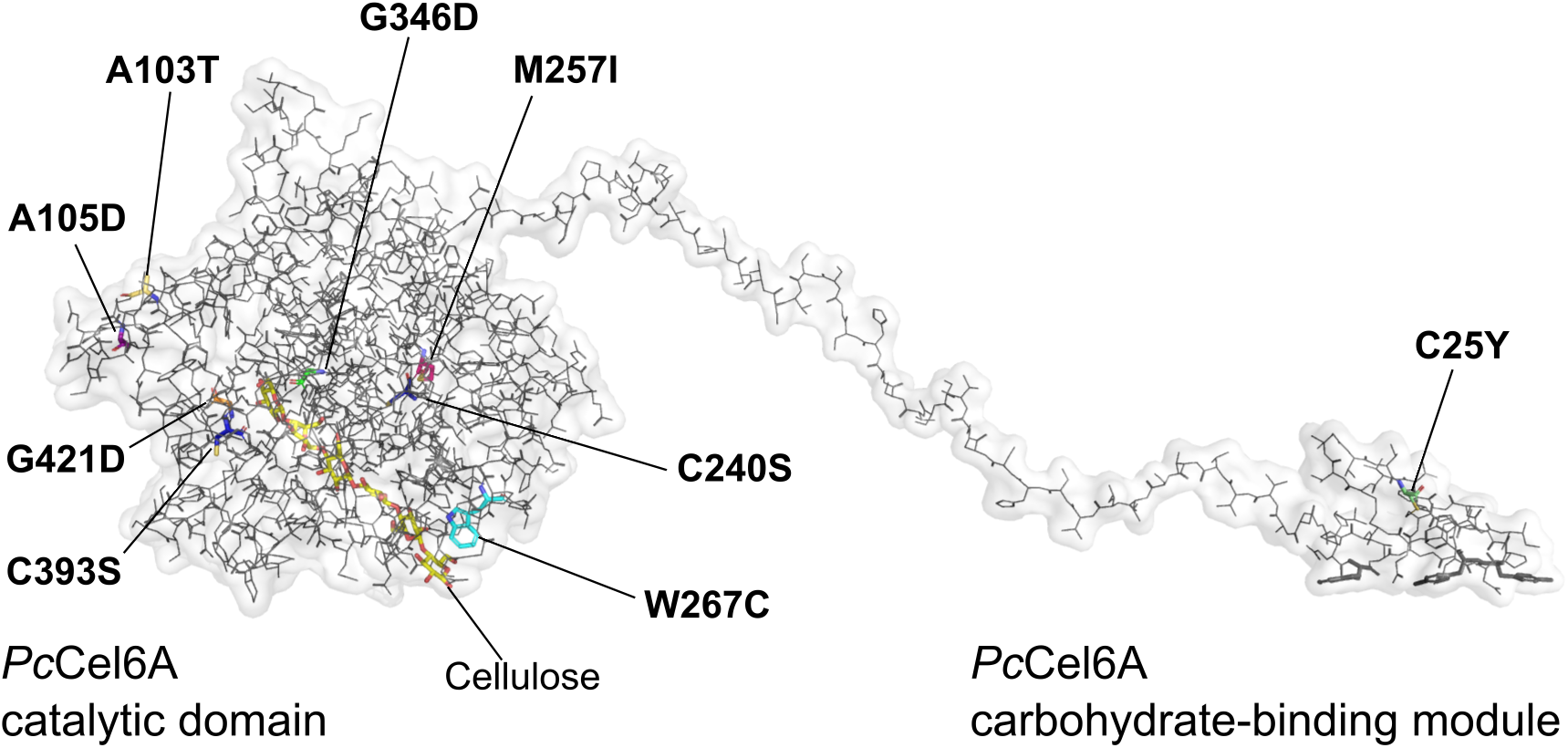
Residues of *Pc*Cel6A selected for mutagenesis. Mutations C25Y, A103T, A105D, M257I, W267C, G346D, and G421D were selected on the basis of a random mutagenesis experiment (see the text). G421A was chosen to test the effect of removing Gly421. C240S/C393S were selected by rational design (see the text). The overall structure of *Pc*Cel6A was created by superposing the structure of the catalytic domain was taken from PDB ID 5XCY,^14)^ and the structure of the carbohydrate-binding module was predicted using Protein Homology/analogy Recognition Engine V 2.0 (Phyre^2^) on the overall structure of *Tr*Cel6A modelled previously. ^18)^ The points of mutations were introduced into images of these domains with the PyMOL Molecular Graphics System, Version 1.0.0.0 Schrödinger, LLC.

### Expression amount of *Pc*Cel6A mutants

The total protein concentration expressed by transformed *P. pastoris* increased with cultivation time after the induction of protein expression by adding methanol to the culture, as shown in Fig. 2. The production levels of W267C and G421D were higher than that of WT. The content of *Pc*Cel6A analyzed by SDS-PAGE did not reflect the protein concentration, presumably because of differences in glycosylation of the recombinant proteins. Therefore, we treated the samples with endoglycosylase H and α-mannnosidase to remove *N*-glycosylation at Asn398. As shown in Fig. 3, after the deglycosylation procedure, several bands were seen between 37 and 75 kDa. Since the molecular weight of *Pc*Cel6A calculated from the amino acid sequence is 46 kDa, these bands at various molecular weights presumably reflect glycosylation at other sites. Indeed, 25 and 8 Ser/Thr residues in the linker and CD, respectively, are predicted to be *O*-glycosylated by the NetOGlyc 4.0 Server (Ver. 4. 0. 0. 13),^16)^ and it has been shown that the major hydrolysis products of *O*- and *N*-linked saccharides attached to a recombinant protein expressed in *P. pastoris* were *O*-linked dimeric to pentameric oligosaccharides.^17)^ Hence, the amount of *O*-glycosylation can be estimated to correspond to 11 to 25 kDa, which is consistent with the idea that the difference in molecular weight of approximately 10 to 20 kDa between the thickest band of each mutant and the calculated value based on the amino acid sequence is due primarily to *O*-glycosylation. We consider that minor differences in the glycosylation amount among the mutants should have a negligible impact on our findings, because the effects of glycosylation of the CD of Cel6A CBH from *Trichoderma reesei* (*Tr*Cel6A) on the enzyme structure and interactions with a ligand were minimal.^18)^

**Fig. 2.**
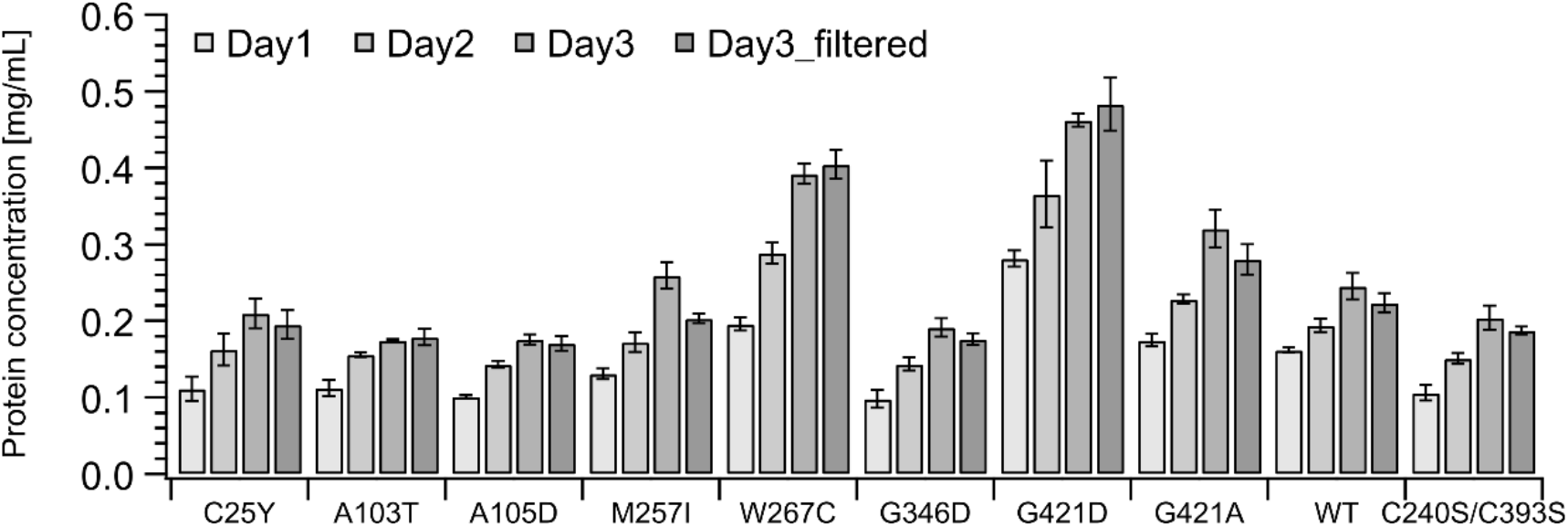
Protein concentration in yeast culture supernatant. Aliquots of 80 μl of yeast culture were sampled for 3 days after induction of the enzyme expression with methanol, and centrifuged at 4 ^°^C with 10,000 x *g* for 10 min. The third-day culture was centrifuged at 4 ^°^C with 3,000 x *g* for 5 min and centrifuged again at 4 ^°^C with 15,000 x *g* for 10 min. The protein concentration of the supernatants was determined using Bio-Rad Protein Assay Dye Reagent Concentrate (Bio-rad Laboratories, Inc.). The absorbance at 595 nm was measured with a Thermo Scientific Multiscan^®^ GO (Thermo Fisher Scientific).

**Fig. 3.**
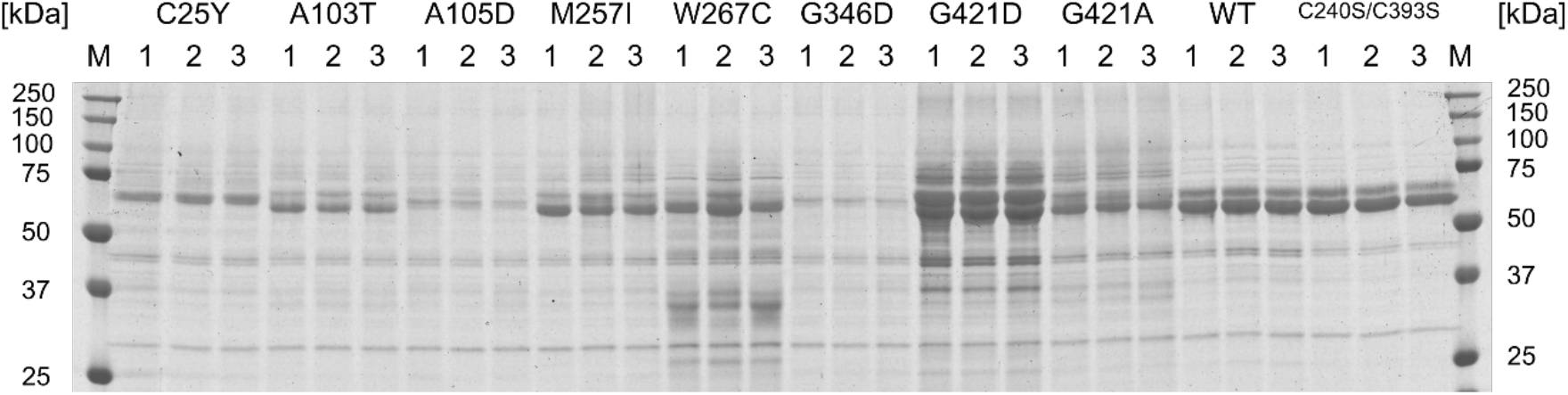
SDS-PAGE of 15 μL *Pc*Cel6A yeast culture supernatants after digestion with endoglycosidase H and α-mannosidase. The gel was imaged with a CanoScan (Canon Inc.), with 50% exposure time. The image was modified to change the shape of the lanes from trapezoidal to rectangular by using GIMP (Ver. 2.10.14.0, https://www.gimp.org) and converted to a 32-bit gray scale with ImageJ. The amount of *Pc*Cel6A was estimated from the peak area of bands between 50 and 75 kDa using ImageJ.

### Thermal properties of *Pc*Cel6A mutants

Hydrolytic activity in the culture supernatant of WT and mutant enzymes was tested using PASC and cellulose III_I_ as substrates. As shown in Fig. 4A, specific activities of PASC hydrolysis by A103T, M257I, and C240S/C393S were significantly improved, compared with WT, at 60 °C. However, when crystalline cellulose III_I_ was used as a substrate, thermal inactivation of A103T, M257I, and WT apparently occurred at 60 °C, though the C240S/C393S double mutant remained active. Ala103 has an α-helix at the surface of the enzyme. It was reported that many advantageous mutations of *Pc*Cel6A based on the sequences of thermophilic fungal Cel6 CBHs are located on the surface of the enzyme.^15)^ Since A103T involves the substitution of compact Ala with bulky Thr, we speculate that its stability might be increased as a result of filling a cavity on the surface of the enzyme that would otherwise be accessible to the solvent. In a previous study, M257I showed an increase of 1.2 ^°^C in the temperature required to reduce the initial activity by 50% within 120 min.^15)^ This might be due to the substitution with a more hydrophobic amino acid, Ile, because Met257 is surrounded by hydrophobic side chains of amino acids in the α-helix.^14)^ If we compare the amino acid sequences of the characterized fungal cellobiohydrolase Cel6s, branched chain amino acids such as Leu that are more hydrophobic than Met appear frequently at the position corresponding to Met257 in *Pc*Cel6A. Thus, it is plausible that these hydrophobic residues improve the stability of the enzyme, since they are highly conserved among fungal Cel6 CBHs.

**Fig. 4.**
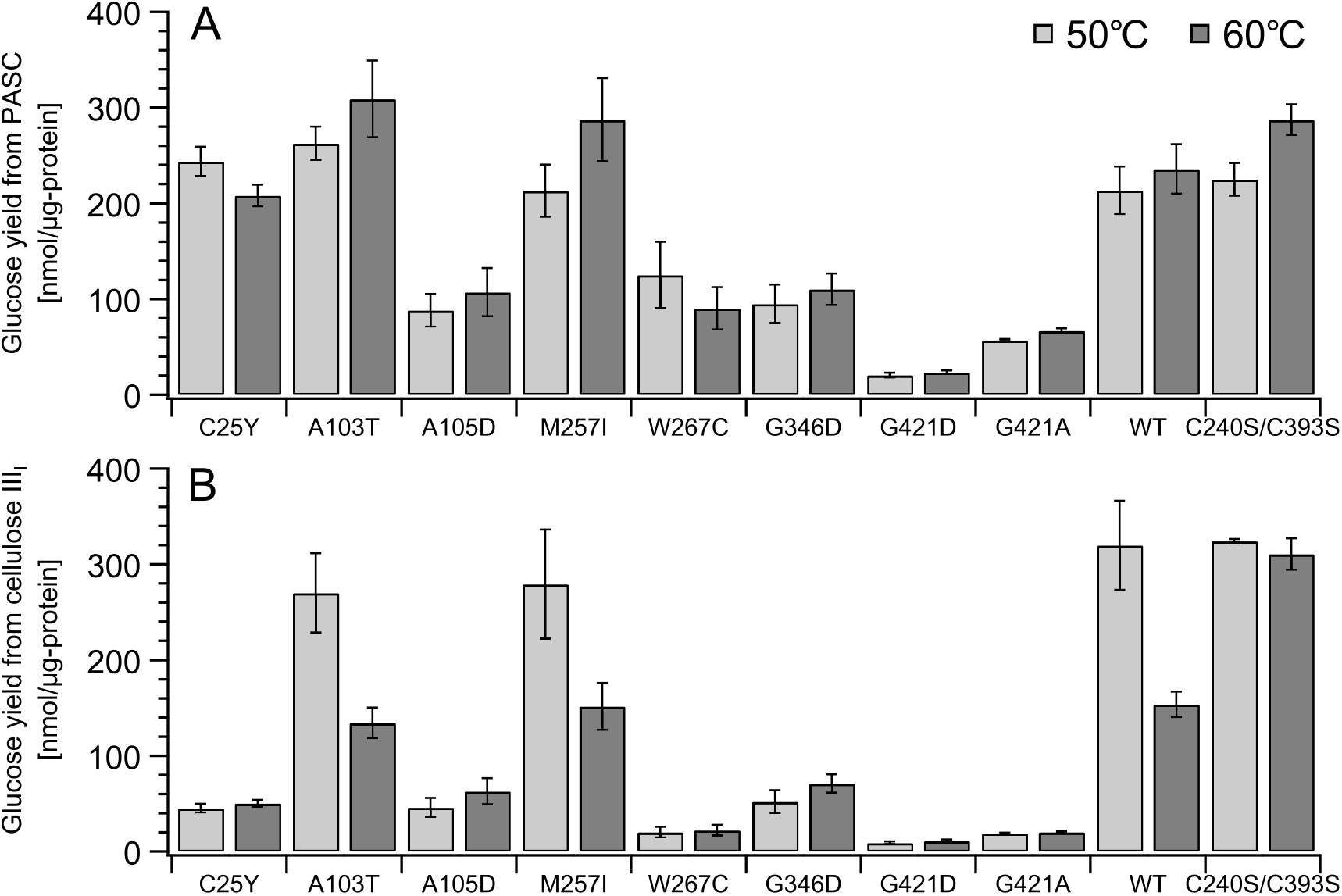
Specific hydrolytic activities towards PASC (A) and cellulose III_I_ (B). *Pc*Cel6A mutants and WT in yeast culture supernatants were incubated at 50 and 60 ^°^C with 0.05% PASC or cellulose III_I_. N = 3. Error bars represent ± 1 standard deviation.

Comparison of the hydrolytic activities towards amorphous (PASC) versus crystalline cellulose (cellulose III_I_) is helpful to judge the origin of activity changes. When the activity is plotted by putting the glucose yield from PASC on the horizontal axis and that from cellulose III_I_ on the vertical axis, we typically see second-degree polynomial curves, as we previously reported.^9)^ The plot should deviate from the curve if the relative activity, i.e., amorphous vs. crystalline, is changed by mutation. As shown in Fig. 5, the activity of C25Y and W267C towards crystalline cellulose was lower at 50 ^°^C (Fig. 5A). Cys25 and Trp267 are located in the CBM and at the entrance to the catalytic tunnel, respectively. Since Cys25 forms a disulfide bond with Cys8 in the CBM, and the replacement of Cys25 with Tyr hinders disulfide bond formation in the CBM, C25Y might reduce the affinity for the surface of crystalline cellulose. Trp267 is located at the entrance of the active site tunnel and plays an important role in the degradation of crystalline cellulose by guiding a cellulose chain into the tunnel, as demonstrated in the case of *Tr*Cel6A.^19)^ Therefore, it seems likely that W267C was unable to take a cellulose chain into the active site tunnel efficiently following the substitution of the nonpolar amino acid Trp with the polar amino acid Cys. Moreover, the substitution of Trp267 with Cys introduces an additional free cysteine, which might negatively affect the expression or the stability of the mutant (Fig. 3). As shown in Fig. 5C and 5D, the division of the glucose production by the protein amount in the reaction solution enables us to compare the catalytic efficiency itself, because it eliminates the effect of the expression level. Although A103T, M257I, and WT showed similar ratios of amorphous and crystalline cellulose degradation to C240S/C393S at 50^°^C (Fig. 5C), they showed reduced ability to degrade crystalline cellulose at 60 ^°^C (Fig. 5D). The mechanism of this effect will be discussed later.

**Fig. 5.**
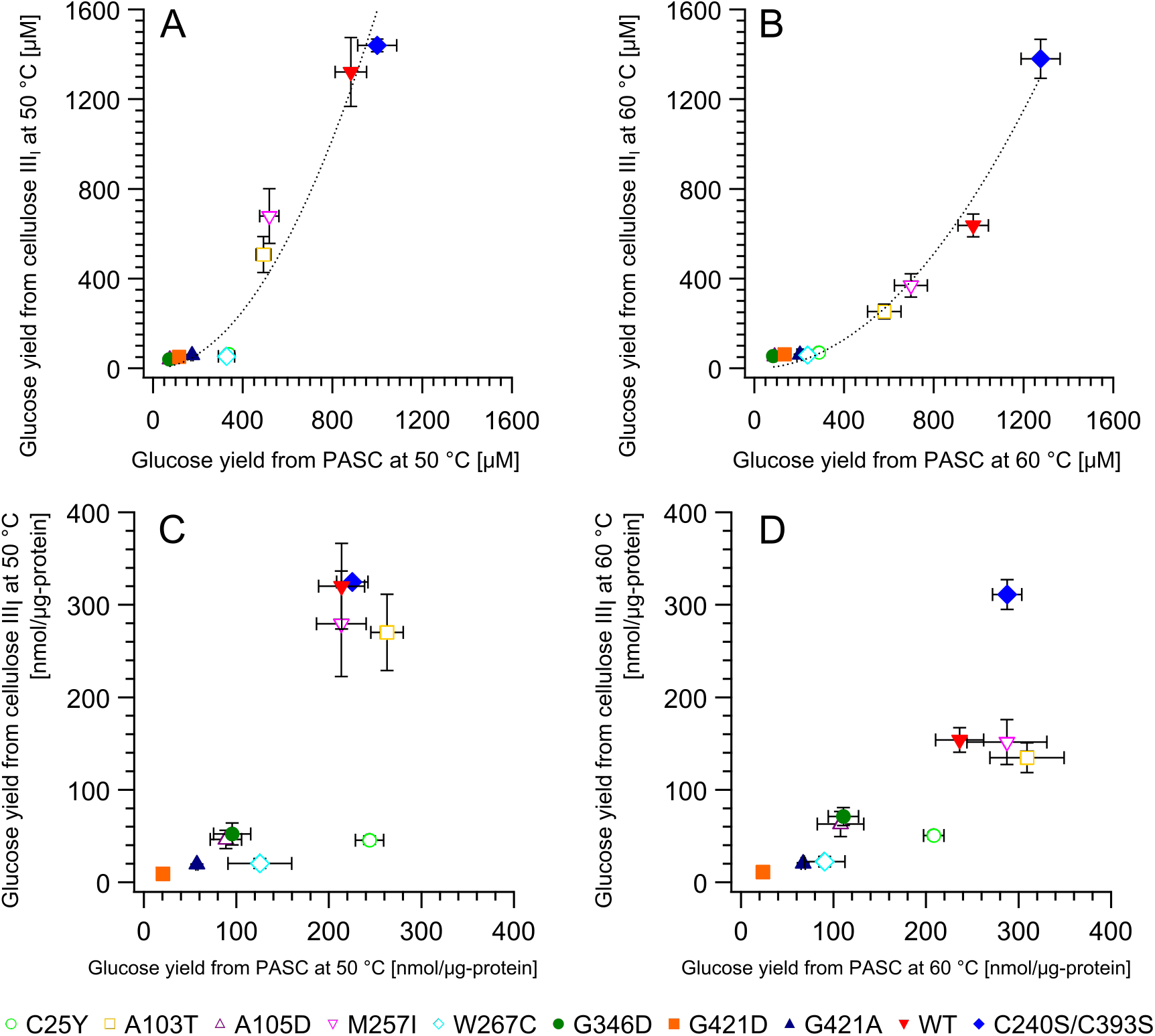
Hydrolytic activities towards PASC and cellulose III_I_ at 50 ^°^C (A) and 60 ^°^C (B), and specific activity at 50 ^°^C (C) and 60 ^°^C (D). The reaction conditions were the same as for Fig. 4. N = 3. Error bars represent ± 1 standard deviation.

When the hydrolytic activity in the reaction at 50 ^°^C is plotted on the horizontal axis against that at 60 ^°^C on the vertical axis, most mutants lie on the same line, as shown in Fig. 6A and 6B. This regression line should reflect the increase in the activity due to the larger kinetic energy generated by raising the temperature in competition with the decrease in the activity due to thermal inactivation of the enzymes. However, C240S/C393S lies far above the line of WT (Fig. 6B), indicating that the double mutant really is a thermostable enzyme favorable for the degradation of amorphous and crystalline cellulose. It has been suggested that disulfide bond cleavage and thiol-disulfide exchange are involved in the thermal inactivation of fungal Cel6 CBHs, because the mutants lacking free cysteine retain their activity to a certain extent even after incubation for 15 min at 90 ^°^C, while the parent CBHs completely loses activity under these conditions.^11)^ Oxidation of thiol might also have a negative effect.^20)^

**Fig. 6.**
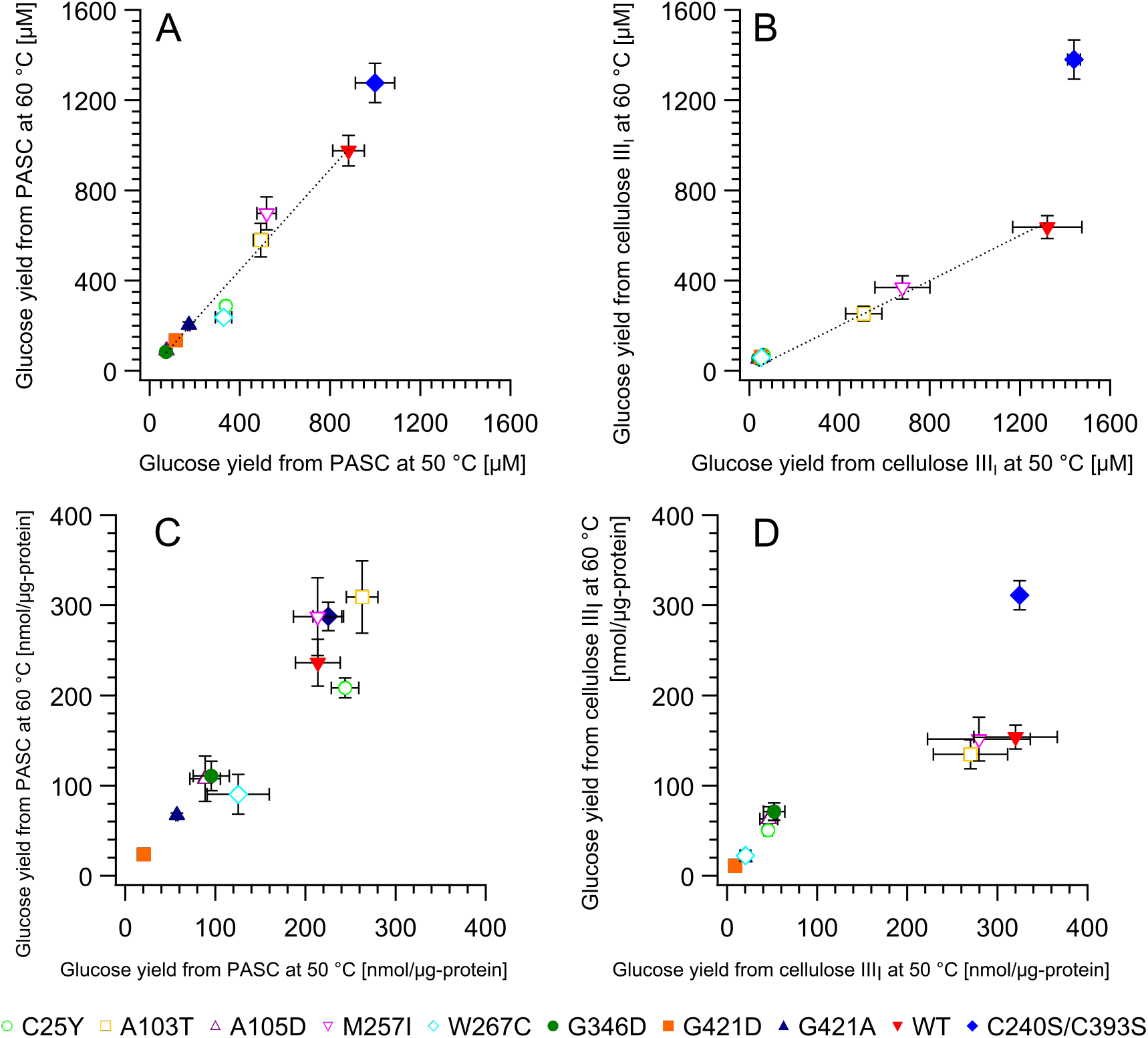
Comparison of the hydrolytic activities at 50 ^°^C and 60 ^°^C towards PASC (A) and cellulose III_I_ (B), and specific activities towards PASC (C) and cellulose III_I_ (D). The reaction conditions were the same as for Fig. 4. N = 3. Error bars represent ± 1 standard deviation.

These effects of free cysteine might explain the decline in the specific activity of A103T, M257I, and WT towards crystalline cellulose at 60 ^°^C (Fig. 5D and 6D). As shown in Fig. 7, Gln187 and Cys408 that are supposed to be directly or indirectly interacting with Cys240 and Cys393, respectively, take double conformation. Therefore, thermal stabilization of C240S/C393S might be resulted from stabilizing these double conformational residues. Simulations suggest that processive cellobiohydrolases are more likely to perform the rate-limiting step of dissociation from crystalline cellulose by backing up along the cellulose chain without opening the substrate-enclosing loops rather than by opening the loops.^21)^ The fact that the mobility of the C-terminal loop (amino acid 390-425) is calculated to be less than that of the N-terminal loop (amino acid 174-178) ^14)^ may be related to the existence of a disulfide bond (Cys361-Cys408) near Cys393. If Cys393 weakens this disulfide bond via Asn162 (and perhaps H_2_O), and the bond is consequently cleaved more readily, the C-terminal loop might not retain its immobile structure at higher temperature, which could interfere with dissociation of the enzyme from crystalline cellulose. Moreover, since the number of free cysteine residues is 0 in C240S/C393S, 3 in C25Y and W267C, and 2 in other mutants and WT, the specific activity towards amorphous cellulose at the higher temperature (Fig. 6C) might also be connected to the number of free cysteine residues in the enzyme.

**Fig. 7.**
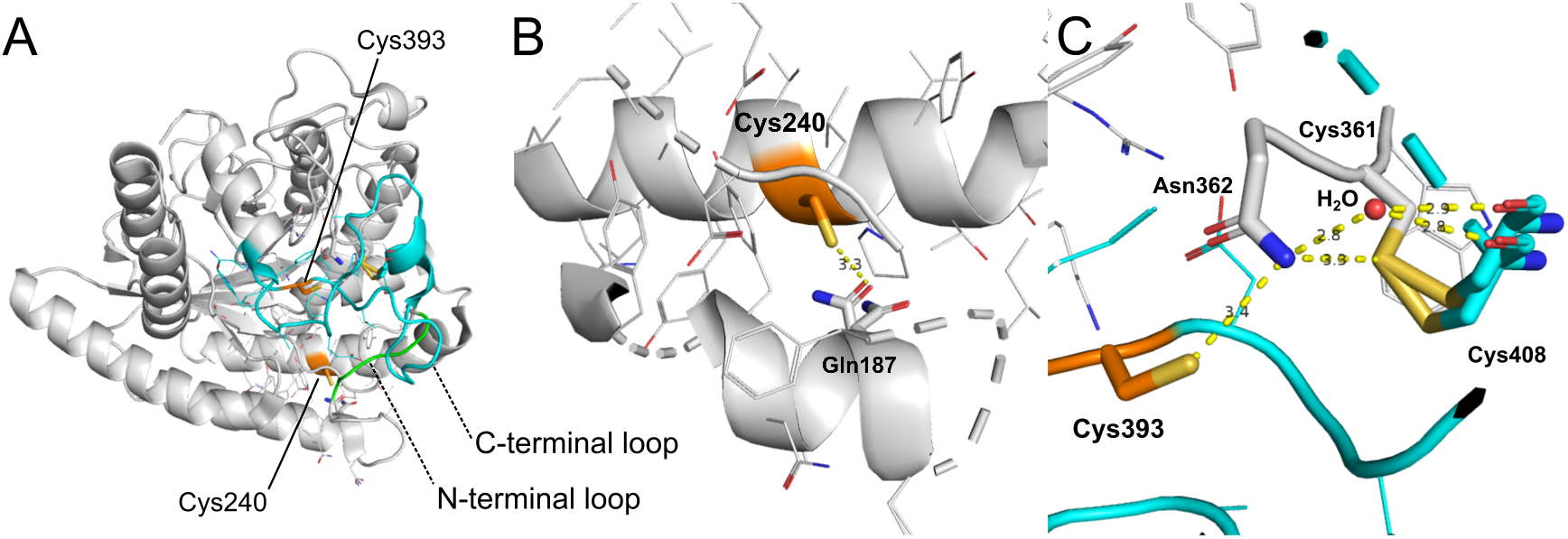
Location of the N- and C- terminal loops (A) and the close-up views of Cys240 (B) and Cys393 (C). The structure of the catalytic domain of *Pc*Cel6A WT was taken from PDB ID 5XCY ^14)^ and modified with the PyMOL Molecular Graphics System. N- and C- terminal loops consist of amino acid residues 174-178 (green) and 390-425 (cyan), respectively. Free cysteine Cys240 and Cys393 are colored orange and shown by sticks. Side chains of the residues around 8 Å from Cys240 and Cys393 are shown by lines. Residues that might be interacting with Cys240 and Cys393 directly or indirectly are (Gln187, Asn362, Cys361 and Cys408) are also represented by sticks.

## CONCLUSION

In this work, we identified several mutants of *Pc*Cel6A that show higher activities than WT at 60 ^°^C. Our results indicate that the number and position of free cysteine residues are critical factors affecting the thermal stability of *Pc*Cel6A. We are currently conducting X-ray crystal structure analysis to better understand the structural basis of the thermal stabilization by the specific substitutions without depending on structure modeling. Our findings should be helpful to increase the efficiency of industrial-scale enzymatic saccharification of cellulose.

## Supporting information

Table S1. Sequence list of primers containing site-directed mutations.

## Abbreviations

CBH: cellobiohydrolase
CBM: carbohydrate-binding module
CD: catalytic domain
GH: glycoside hydrolase
PASC: phosphoric acid-swollen cellulose
WT: wild type

## ACKNOWLEDGEMENTS

S.Y. is grateful for financial support from UTokyo Sustainable Agriculture Education Program during a Master’s course. The authors are grateful for Grants-in-Aid for Scientific Research (B) (15H04526, 18H02252 and 19H03013 to K.I.) from the Japan Society for the Promotion of Science (JSPS), a Grant-in-Aid for Innovative Areas from the Japanese Ministry of Education, Culture, Sports, and Technology (MEXT) (No. 18H05494 to K.I.). In addition, K.I. thanks Business Finland (BF, formerly the Finnish Funding Agency for Innovation (TEKES)) for support via the Finland Distinguished Professor (FiDiPro) Program “Advanced approaches for enzymatic biomass utilization and modification (BioAD)”.

