## Supplementary material for "Thermostable mutants of glycoside hydrolase family 6 cellobiohydrolase from the basidiomycete *Phanerochaete chrysosporium*": Table S1. Sequence list of primers containing site-directed mutations.

**Table S1**. Sequence list of primers containing site-directed mutations. Seven mutations from random mutagenesis experiment except for W267C were designed for the amplification by DNA polymerase PrimeSTAR Max. DNA polymerase KOD-Plus was used for the amplification of W267C. The overlap regions are in bold and the mutated codons are underlined in all sequences.

| Mutation | Primer | Sequence (5' to 3') |
|---|---|---|
| C25Y | Forward | **actacctatacggtt**ctcaatccatac |
| | Reverse | **aaccgtataggtagt**accggagacgca |
| A103T | Forward | **gaggtcactgctgct**gctaagcagatc |
| | Reverse | **agcagcagtgacctc**gttcgcgtagta |
| A105D | Forward | **gccgctgatgctaag**cagatcacggat |
| | Reverse | **cttagcatcagcggc**gacctcgttcgc |
| M257I | Forward | **atgtacattgatgct**ggccacgccggc |
| | Reverse | **agcatcaatgtacat**gtacacgccgac |
| G346D | Forward | **gaccaggatcgctcc**ggtgtgcaaaac |
| | Reverse | **ggagcgatcctggtc**gacgatgaaggt |
| G421D | Forward | **gaggccgatacctgg**ttccaggcgtac |
| | Reverse | **ccaggtatcggcctc**aggagcgggctg |
| G421A | Forward | **gaggccgctacctgg**ttccaggcgtac |
| | Reverse | **ccaggtagcggcctc**aggagcgggctg |
| W267C | Forward | cggctgtcccgcgaacctgtcg |
| | Reverse | agccagccggcgtggccagc |
